# Postnatal maturation of *in vivo* membrane potential dynamics during hippocampal ripples

**DOI:** 10.1101/2023.02.10.527963

**Authors:** Asako Noguchi, Yuji Ikegaya

## Abstract

Sharp-wave ripples (SWRs) are transient high-frequency oscillations of local field potentials (LFPs) in the hippocampus and play a critical role in memory consolidation. During SWRs, CA1 pyramidal cells exhibit rapid spike sequences that often replay the sequential activity that occurred during behavior. This temporally organized firing activity gradually emerges during two weeks after the eye opening; however, it remains unclear how the organized spikes during SWRs mature at the intracellular membrane potential (*V*m) level. Here, we recorded the *V*m of CA1 pyramidal cells simultaneously with hippocampal LFPs from immature mice after the developmental emergence of SWRs. On postnatal days 16–17, *V*m dynamics around SWRs were premature, characterized by prolonged depolarizations without either pre- or post-SWR hyperpolarization. The biphasic hyperpolarizations, which are a typical feature of adult SWR-relevant *V*m, formed by approximately postnatal day 30. This *V*m maturation was associated with an increase in SWR-associated inhibitory inputs to pyramidal cells. Thus, the development of SWR-relevant inhibition restricts the temporal windows for spikes of pyramidal cells and allows CA1 pyramidal cells to organize their spike sequences during SWRs.

**Significance statement:** Sharp-wave ripples (SWRs) are prominent hippocampal oscillations and play a critical role in memory consolidation. During SWRs, hippocampal neurons synchronously emit spikes with organized temporal patterns. This temporal structure of spikes during SWRs develops during the third and fourth postnatal weeks, but the underlying mechanisms are not well-understood. Here, we recorded *in vivo* membrane potentials from hippocampal neurons in premature mice and suggest that the maturation of SWR-associated inhibition enables hippocampal neurons to produce precisely controlled spike times during SWRs.

## Introduction

The hippocampus is a critical brain region for representing the outer world, internalizing behavioral trajectories of animals into their mental maps (O’Keefe and Nadel, 1979; Ekstrom and Ranganath, 2018), and consolidating the acquired internal information into memory episodes (Girardeau and Zugaro, 2011). During quiet wakefulness and slow wave sleep, the hippocampus generates sharp-wave ripples (SWRs), which are brief (40– 100 ms) and high frequency (100–250 Hz) oscillations in the local field potentials (LFPs) (Buzsáki, 2015). SWRs are associated with spikes emitted sequentially by multiple CA1 pyramidal cells, which often reflect offline reactivation of spike sequences that occurred during behavior and are believed to contribute to memory consolidation (Lee and Wilson, 2002; Girardeau and Zugaro, 2011; Foster, 2017). A recent study demonstrated that these temporally organized spikes emerged gradually during the third and fourth postnatal weeks; sequential spike patterns appeared in the third postnatal week and matured enough to represent behavioral trajectories during the fourth postnatal week (Farooq and Dragoi, 2019). However, it is still unclear how the maturation of temporally coordinated spikes during SWRs is implemented at the single-cell level.

In adult rodents, SWR-associated spikes of CA1 pyramidal cells are regulated at the subthreshold level. Pyramidal cells exhibit triphasic membrane potential (*V*m) dynamics, *i*.*e*., pre-SWR hyperpolarization, depolarization during SWRs, and post-SWR hyperpolarization (English et al., 2014; Hulse et al., 2016; Noguchi et al., 2022). These *V*m dynamics coordinate the characteristic inhibitory and excitatory balance around SWRs and regulate the spike times of individual neurons during SWRs. Extracellular recordings from freely behaving animals reveal that inhibitory interneurons regulate the spike times of pyramidal cells (Stark et al., 2014, 2015; Noguchi et al., 2022). Given that CA1 interneurons are still in the process of maturing during the third and fourth postnatal weeks (Cohen et al., 2000; Doischer et al., 2008; Le Magueresse and Monyer, 2013), the maturation of local inhibitory circuits in the CA1 subregion could be a potential driver behind the postnatal refinement of SWR-related activity. Based on this idea, we hypothesized that premature CA1 pyramidal cells exhibit different SWR-associated *V*m dynamics than mature cells.

In the present study, we recorded intracellular *V*m activity of CA1 pyramidal neurons from mice aged between postnatal day 16 (P16) and P40 and suggested that inhibitory synaptic inputs that increase gradually during development shape the characteristic *V*m dynamics around SWRs, which possibly underlies the maturation of temporally regulated SWR-associated spikes of pyramidal cells during the third and fourth postnatal weeks.

## Materials and Methods

### Animal ethics

Animal experiments were performed with the approval of the animal experiment ethics committee at the University of Tokyo (approval numbers: P4-4) and in accordance with the University of Tokyo guidelines for the care and use of laboratory animals. These experimental protocols were conducted in accordance with the Fundamental Guidelines for the Proper Conduct of Animal Experiments and Related Activities in Academic Research Institutions (Ministry of Education, Culture, Sports, Science and Technology, Notice No. 71 of 2006), the Standards for Breeding and Housing of and Pain Alleviation for Experimental Animals (Ministry of the Environment, Notice No. 88 of 2006) and the Guidelines on the Method of Animal Disposal (Prime Minister’s Office, Notice No. 40 of 1995).

### Patch-clamp and LFP recordings

Whole-cell recordings were obtained from postnatal 16– to 17-, 21– to 22-, 27– to 28-, and 29– to 40-day-old ICR mice (Japan SLC, Shizuoka, Japan) (Ishikawa et al., 2014; Funayama et al., 2015). After exposure to an enriched environment for 30 min, the mice were intraperitoneally anesthetized with 1.5 g/kg (P16–17 and P21–22 mice) or 2.25 g/kg (P27–28 and P29–40 mice) urethane. P16–17 mice were not exposed to an enriched environment because they did not exhibit exploratory behavior at this developmental stage. Anesthesia was confirmed by the absence of paw withdrawal, whisker movement, and eyeblink reflexes. The skin was subsequently removed from the head, and a metal head-holding plate was fixed to the skull. A craniotomy was performed, the coordinates of which were changed according to the size of the brain at different developmental periods. Based on the length between the bregma and the lambda, the skull was cut into squares with one side five-eighth the length. The anteromedial corner of the square was at one-eighth of the length posterior and lateral to the bregma. The neocortex above the hippocampus was subsequently aspirated (Kuga et al., 2011; Sakaguchi et al., 2012; Matsumoto et al., 2016). The exposed hippocampal window was covered with 2.0% agar at a thickness of 1.5 mm. LFPs were recorded from the dorsal CA1 region using a tungsten electrode (3.5–4.5 MΩ, catalog #UEWMGCSEKNNM, FHC, USA) coated with a crystalline powder of 1,1′-dioctadecyl-3,3,3′,3′-tetramethylindocarbocyanine perchlorate (DiI). Whole-cell patch-clamp recordings were obtained from neurons in the CA1 pyramidal cell layer 0.8-1.2 mm below the hippocampal window using borosilicate glass electrodes (3-8 MΩ). Pyramidal cells were identified based on regular spiking properties in response to step-pulse current injection and morphological features, including apical, oblique, and basal dendrites with spines, in *post hoc* histology experiments. A cell was discarded unless it was identified as a pyramidal cell. For current-clamp recordings, the intrapipette solution consisted of the following reagents: 120 mM K-gluconate, 10 mM KCl, 10 mM HEPES, 10 mM creatine phosphate, 4 mM MgATP, 0.3 mM Na2GTP, 0.2 mM EGTA (pH 7.3), and 0.2% biocytin. Cells were discarded when the mean resting potential exceeded -55 mV and the action potentials were below -20 mV. For voltage-clamp recordings, the intrapipette solution consisted of the following reagents: 130 mM CsMeSO4, 10 mM CsCl, 10 mM HEPES, 10 mM phosphocreatine, 4 mM MgATP, 0.3 mM NaGTP, and 10 mM QX-314. Cells were discarded if the access resistance exceeded 60 MΩ. Signals recorded by LFP electrodes were amplified using a DAM80 AC differential amplifier. Signals recorded by patch-clamp electrodes were amplified using MultiClamp 700B amplifiers. Both types of signals were digitized at a sampling rate of 20 kHz using a Digidata 1440A digitizer that was controlled by pCLAMP 10.3 software (Molecular Devices).

### Histology

Following each experiment, the electrode was carefully withdrawn from the hippocampus. The mice were transcardially perfused with 4% paraformaldehyde followed by overnight postfixation. The brains were coronally or sagittally sectioned at a thickness of 100 μm using a vibratome. The sections were incubated with 2 μg/ml streptavidin-Alexa Fluor 594 conjugate and 0.2% Triton X-100 for 4 h, followed by incubation with 0.4% NeuroTrace 435/455 blue fluorescent Nissl stain (Thermo Fisher Scientific; N21479) for 2-4 h. The location of LFP electrodes was also detectable via DiI fluorescence. Fluorescence images were acquired using an FV1200 (Olympus) or A1 HD25 (Nikon) confocal microscope and were subsequently merged.

### Data analysis

Data were analyzed offline using custom-made MATLAB (R2017b, Natick, Massachusetts, USA) programs. The summarized data are reported as the mean ± SD unless otherwise specified. For box plots, the centerline shows the median, the box limits show the upper and lower quartiles, and the whiskers cover the 10− 90% quantiles. *P* < 0.05 was considered statistically significant. All statistical tests were two-sided.

### SWR analysis

To detect SWRs from LFP traces recorded by a tungsten electrode, LFP traces were downsampled to 500 Hz and bandpass filtered between 100 and 250 Hz. Ripples, referred to here as SWRs, were detected at a threshold of 3 × SD of the baseline noise (Mizunuma et al., 2014). The detected events were subsequently scrutinized by eye and manually rejected if the detection was erroneous. The ripple power and frequency for each SWR event were calculated as the maximum power and its frequency between 100 and 250 Hz after the fast Fourier transform analyses of the raw traces.

### ΔVm analysis

Spikes were detected as peaks during periods with *V*m greater than -20 mV in raw 20-kHz traces of whole-cell recordings. To average subthreshold *V*m values, spikes were truncated as follows: i) raw *V*m traces were smoothed using the moving average with a window of 1 ms, ii) the minimum index at which the rate of change exceeded 4 V/s was detected as the leading edge of a spike, and iii) traces were linearly interpolated from the leading edge until the time point that first showed a *V*m value below the edge value. When multiple edges, such as those of complex spikes, were detected within a period of 30 ms, these edges were individually interpolated from the previous edge until the next time point that showed a *V*m value below the edge value. They were aligned to the SWR onset times and additively averaged across SWRs.

To quantify the full widths at half maximum (FWHM) of depolarizations during SWRs, we detected the highest peak value of *V*m between -20 ms and +120 ms relative to the SWR onset time as the maximum. As the baseline *V*m, *V*m between -2,000 ms and -1,000 ms relative to the SWR onset time was averaged. For the SWR events in which the maximum was above the baseline *V*m values, FWHM was calculated. To quantify Δ*V*m_pre_ and Δ*V*m_post_, we identified the time point that gave the minimum *V*m between -50 and 0 ms or 100 and 200 ms relative to the SWR onset time (pre-SWR and post-SWR period, respectively). *V*m was averaged between -25 ms and +25 ms relative to these time points, and the baseline *V*m was subtracted to obtain Δ*V*m_pre_ and Δ*V*m_post_, respectively. Δ*V*m_during_ was obtained by subtracting the baseline *V*m value from the maximum *V*m value between 0 and 100 ms relative to the SWR onset time (during-SWR period).

For SWR-triggered analysis of excitatory and inhibitory postsynaptic conductances (EPSGs and IPSGs), we took a postsynaptic current at a given time between -2,000 ms and +400 ms relative to the SWR onset and subtracted the mean current between -2,000 ms and -1,000 ms. The current traces were averaged across SWRs. EPSGs and IPSGs were calculated as *I*_E/I_/(*V*_h_ − *E*_rev, E/I_), where *I*_E/I_ is the excitatory/inhibitory current at a given time, *V*h is the holding potential of -70 mV and +10 mV for EPSGs and IPSGs, respectively (Gan et al., 2017), and *E*_rev, E/I_ are the reversal potentials of 0 mV and -90 mV for EPSGs and IPSGs, respectively (Funayama et al., 2016).

## Results

### Higher membrane excitability and fewer spike bursts in premature hippocampal neurons

To examine the developmental changes in SWR-associated *V*m dynamics of hippocampal neurons, we conducted *in vivo* whole-cell patch-clamp recordings from dorsal hippocampal CA1 pyramidal cells simultaneously with LFP recordings in the CA1 region of premature mice. Recordings were excluded from subsequent analyses if either *post hoc* biocytin-based visualization, their recording sites, or their firing properties failed to identify the recorded neurons as CA1 pyramidal cells (Noguchi et al., 2022). SWRs start to emerge after the end of the second postnatal week (Buhl and Buzsáki, 2005). Indeed, we did not detect SWRs in mice younger than P15 (*n* = 6 mice) based on the same detection criteria used in adult mice. Thus, we collected data from mice older than P15. As a result, we recorded *V*ms from 21, 14, 9 and 44 patch-clamped cells in a total of 15, 7, 5 and 20 mice aged P16–17, P21–22, P27–28 and P29–40, respectively. Recording periods ranged from 162 s to 2749 s (median = 586 s), 147 s to 2116 s (median = 890 s), 420 s to 1380 s (median = 496 s) and 113 s to 2,097 s (median = 546 s), during which a total of 240, 664, 385, and 1882 SWRs were detected in LFPs, respectively.

We first compared the subthreshold *V*m recorded from premature (P16–17) and adult (P29–40) mice, regardless of SWR occurrence (Figure 1A, B). P16–17 mice showed significantly higher firing rates than P29–40 mice, indicating higher excitability in immature pyramidal cells (Figure 1C, *P* = 0.041, *t*_63_ = 2.1, Student’s *t*-test, *n* = 21 and 44 cells for P16–17 and P29–40, respectively). Consistently, the mean *V*m values were significantly higher in P16–17 mice than in P29–40 mice (Figure 1D, *P* = 0.0028, *t*_63_ = 3.1, Student’s *t-*test, *n* = 21 and 44 cells for P16–17 and P29–40, respectively). The fluctuations in *V*m, captured by their standard deviations (SDs), did not differ between premature and adult mice (Figure 1E, *P* = 0.22, *t*_63_ = 1.3, Student’s *t*-test, *n* = 21 and 44 cells for P16–17 and P29–40, respectively). Notably, burst-like firing activity was observed in both developmental periods (Figure 1A, B bottom), but the interspike intervals (ISIs) in the burst-like activity in P16–17 mice were larger than those in P29– 40 mice (Figure 1F, *P* = 3.3×10^−231^, *D* = 0.50, two-sample Kolmogorov‒Smirnov test for the cumulative distributions of ISIs less than 200 ms, *n* = 3269 and 1539 intervals for P16–17 and P29–40, respectively). Smaller ISIs on P16–17 suggest less burst-like firing activity in immature mice. We defined a series of successive spikes with all ISIs less than 8 ms as a burst event and calculated the frequency of burst events for each cell (Harris et al., 2001; Mizuseki and Buzsáki, 2013). The burst event rates were significantly lower in P16–17 mice than in P29–40 mice (Figure 1F inset, *P* = 0.025, *t*_63_ = -2.3, Student’s *t*-test, *n* = 21 and 44 cells for P16–17 and P29–40, respectively). These results indicate that at the beginning of SWR emergence, hippocampal CA1 pyramidal neurons exhibit high excitability with higher spontaneous spike rates but rarely fire spikes at frequencies higher than ∼100 Hz.

**Figure 1.**
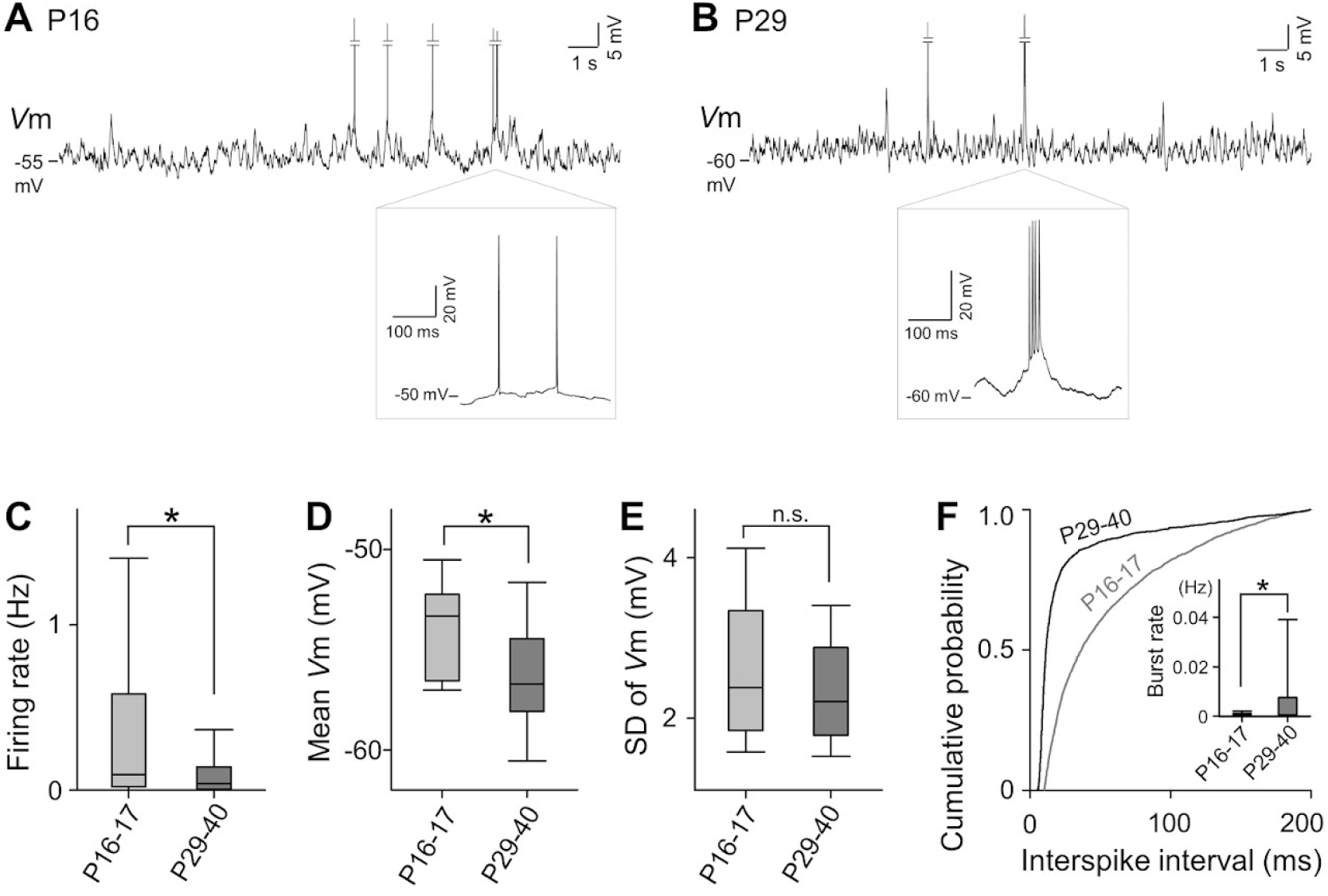
Comparison of *in vivo* spiking and *V*m dynamics of hippocampal CA1 pyramidal cells between P16–17 and P29–40 mice. (**A**) A representative *V*m trace of a hippocampal CA1 pyramidal neuron recorded from a P16 mouse, a part of which is enlarged in the bottom box. (**B**) The same as A, but from a P29 mouse. (**C**) Firing rates were higher in P16–17 mice than in P29–40 mice. *P* = 0.041, *t*_63_ = 2.1, Student’s *t*-test, *n* = 21 and 44 cells for P16–17 and P29–40 mice, respectively. (**D**) Mean *V*ms were higher in P16–17 mice than in P29–40 mice. *P* = 0.0028, *t*_63_ = 3.1, Student’s *t-*test, *n* = 21 and 44 cells for P16–17 and P29–40 mice, respectively. (**E**) The SD values of *V*m were not different between P16–17 and P29–40 mice. *P* = 0.22, *t*_63_ = 1.3, Student’s *t*-test, *n* = 21 and 44 cells for P16–17 and P29–40, respectively. (**F**) Cumulative probability distributions of interspike intervals (ISIs) were different between P16–17 and P29–40 mice. Data are displayed in the range of 200 ms or less. *P* = 3.3×10^−231^, *D* = 0.50, two-sample Kolmogorov‒Smirnov test, *n* = 3269 and 1539 intervals for P16–17 and P29–40, respectively. Inset: The event rates of burst spikes (ISI < 8 ms) were lower in P16–17 mice than in P29–40 mice. *P* = 0.025, *t*_63_ = -2.3, Student’s *t*-test, *n* = 21 and 44 cells for P16–17 and P29–40 mice, respectively.

### Maturation of SWRs throughout the third and fourth postnatal weeks

We investigated the developmental maturation of SWR events. As shown in the representative LFP traces in Figure 2A and B, the SWR event frequency was higher in older mice. Data are summarized in Figure 2C, indicating a significant increase during the fourth postnatal week (Figure 2C, \**P* < 0.05, one-way analysis of variance (ANOVA) followed by Tukey‒Kramer *post hoc* test, *n* = 21, 14, 5 and 20 datasets for P16–17, P21– 22, P27–28 and P29–40, respectively). Similar results were obtained for ripple power and frequency but not for SWR duration (Figure 2D-F, \**P* < 0.05, one-way analysis of variance (ANOVA) followed by Tukey‒Kramer *post hoc* test, *n* = 240, 664, 385 and 1882 SWRs for P16–17, P21–22, P27–28 and P29–40, respectively). In addition to the change in the characteristics of individual SWRs, we noticed that successive SWRs (Davidson et al., 2009; Yamamoto and Tonegawa, 2017; Oliva et al., 2018) occurred more frequently in adult mice than in P16–17 mice (Figure 2A, B).

**Figure 2.**
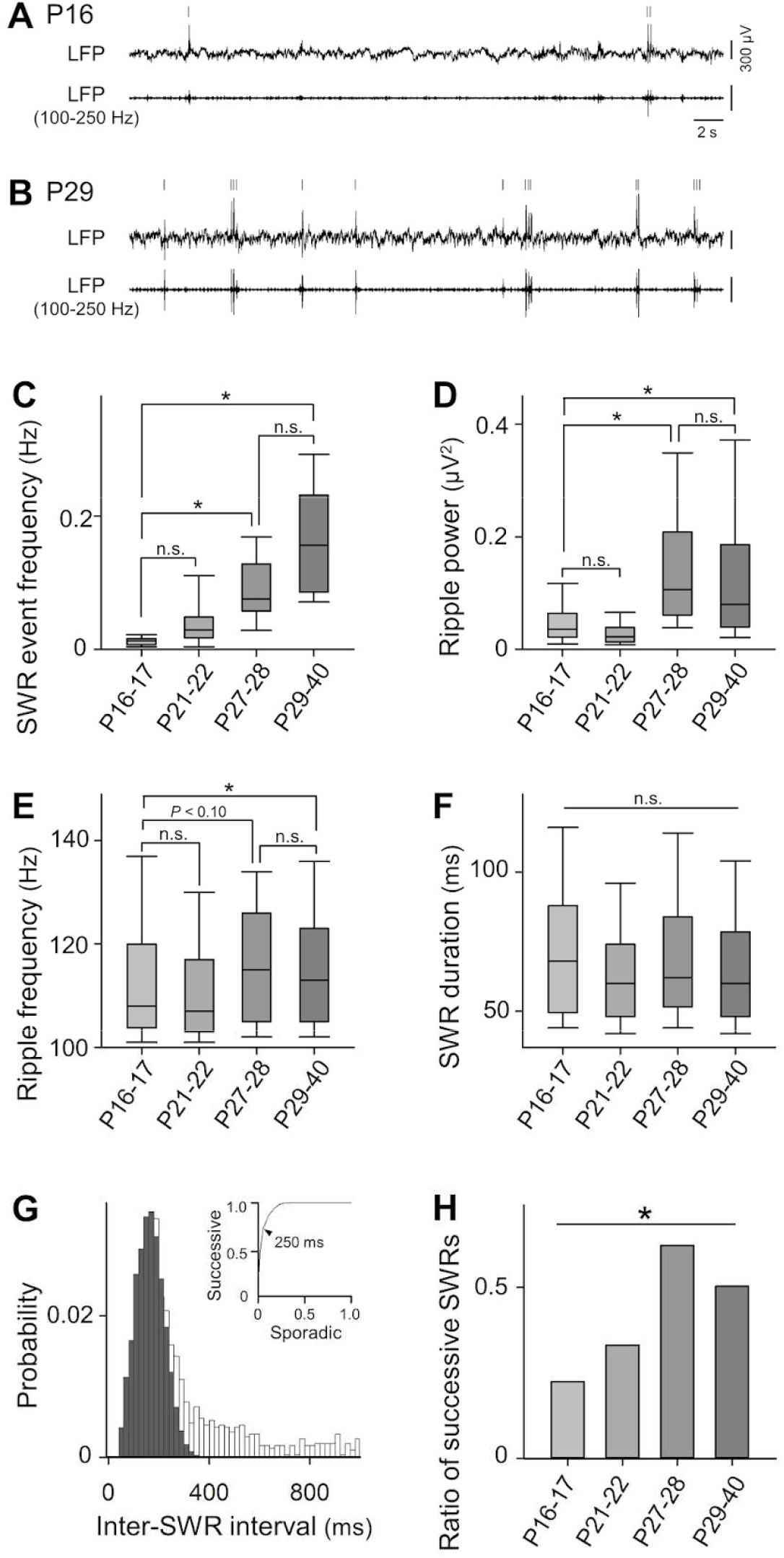
Developmental changes in SWR characteristics. (**A**) A representative raw trace of hippocampal LFPs recorded from a P16 mouse (top) that were bandpass-filtered between 100 and 250 Hz (bottom). Ticks above the traces indicate the onset times of SWRs. (**B**) The same as A, but from a P29 mouse. (**C**) The mean frequency of SWR events increased as postnatal days increased. \**P* < 0.05, Tukey‒Kramer *post hoc* test following one-way analysis of variance (ANOVA), *n* = 21, 14, 5 and 20 datasets from P16–17, P21–22, P27–28 and P29–40 mice, respectively. (**D, E, F**) The same as C, but for the power (D) and frequency (E) of ripples and the duration of SWRs. *n* = 240, 664, 385 and 1882 SWRs from 15, 7, 5 and 20 mice for P16–17, P21–22, P27–28 and P29– 40, respectively. (**G**) Probability distribution of inter-SWR intervals. The distribution of values smaller than the distribution peak (mode) was fitted with a half Gaussian distribution and folded against the mode (black). The distribution in white is the remaining after subtracting the estimated Gaussian (black) distribution. Then, the distribution of inter-SWR intervals was divided into two distributions for successive SWRs (black) and sporadic SWRs (white). Inset: ROC curve with the probability of repetitive SWR intervals as the vertical axis and the probability of sporadic SWR intervals as the horizontal axis. The optimal cutoff was found to be 250 ms, which was set to the threshold of SWR bursts in this study. (**H**) Developmental changes in the ratios of successive SWRs to the total SWRs. *P* = 1.9×10^−12^, χ^2^ = 57.6, Chi-squared test.

To quantitatively define successive SWRs, we sought the upper threshold of interevent intervals (IEIs) based on the pooled distribution of IEIs for all recorded SWR events (Figure 2G). Specifically, the distribution of IEIs less than the mode was fitted with a half Gaussian distribution model and folded around the mode value (Figure 2G, black), which was designated as a population of successive SWRs. The remaining distribution after subtracting this distribution from the entire distribution was considered sporadic SWRs (Figure 2G, white). The receiver operating characteristic curve was drawn based on the ratios of the two distributions to the entire distribution of IEIs less than 1 s, and the threshold IEI for detecting successive SWRs was set to 250 ms, with which the two distributions were best separated (Figure 2G, inset). The ratios of successive SWRs to total SWRs were higher in older mice (Figure 2H, *P* = 1.9×10^−12^, χ^2^ = 57.6, Chi-squared test). Thus, SWRs gradually matured through the third and fourth postnatal weeks.

### Weak SWR selectivity of spikes in premature hippocampal neurons

We compared SWR-associated firing activity of CA1 pyramidal neurons between P16– 17 and P29–40 (Figure 3A). In contrast to the overall firing rates (Figure 1C), the mean firing rates in pre-SWR periods (0–50 ms before SWR onsets) and during-SWR periods (0–100 ms after SWR onsets) were significantly lower in P16–17 mice than in P29–40 mice, while the mean firing rate in post-SWR periods (100–200 ms after SWR onsets) did not differ by age (Figure 3B, *P* = 0.033, 0.0053, and 0.16, *t*_63_ = -2.2, -2.9, and -1.4, Student’s *t*-test, for pre-, during-, post-SWR periods, *n* = 21 and 44 for P16–17 and P29– 40, respectively). Thus, mature pyramidal cells were likely to fire spikes more selectively during SWRs. Consistent with this idea, the distribution of spike latencies relative to the SWR onsets exhibited a sharper peak, and its kurtosis was larger in P29–40 mice than in P16–17 mice (Figure 3C left). To statistically assess the difference in the two kurtosis values (ΔKurtosis), all spike latencies were pooled from P16–17 and P29–40 mice and were randomly reassigned to two groups with the same number of values as the input data. Then, ΔKurtosis was calculated for the resampled groups. This procedure was repeated 1000 times to generate a null distribution. The real ΔKurtosis value was larger than the maximum of the null distribution (*P* < 10^−3^), indicating that spikes were temporally more locked to SWRs on P29–40 than P16–17.

**Figure 3.**
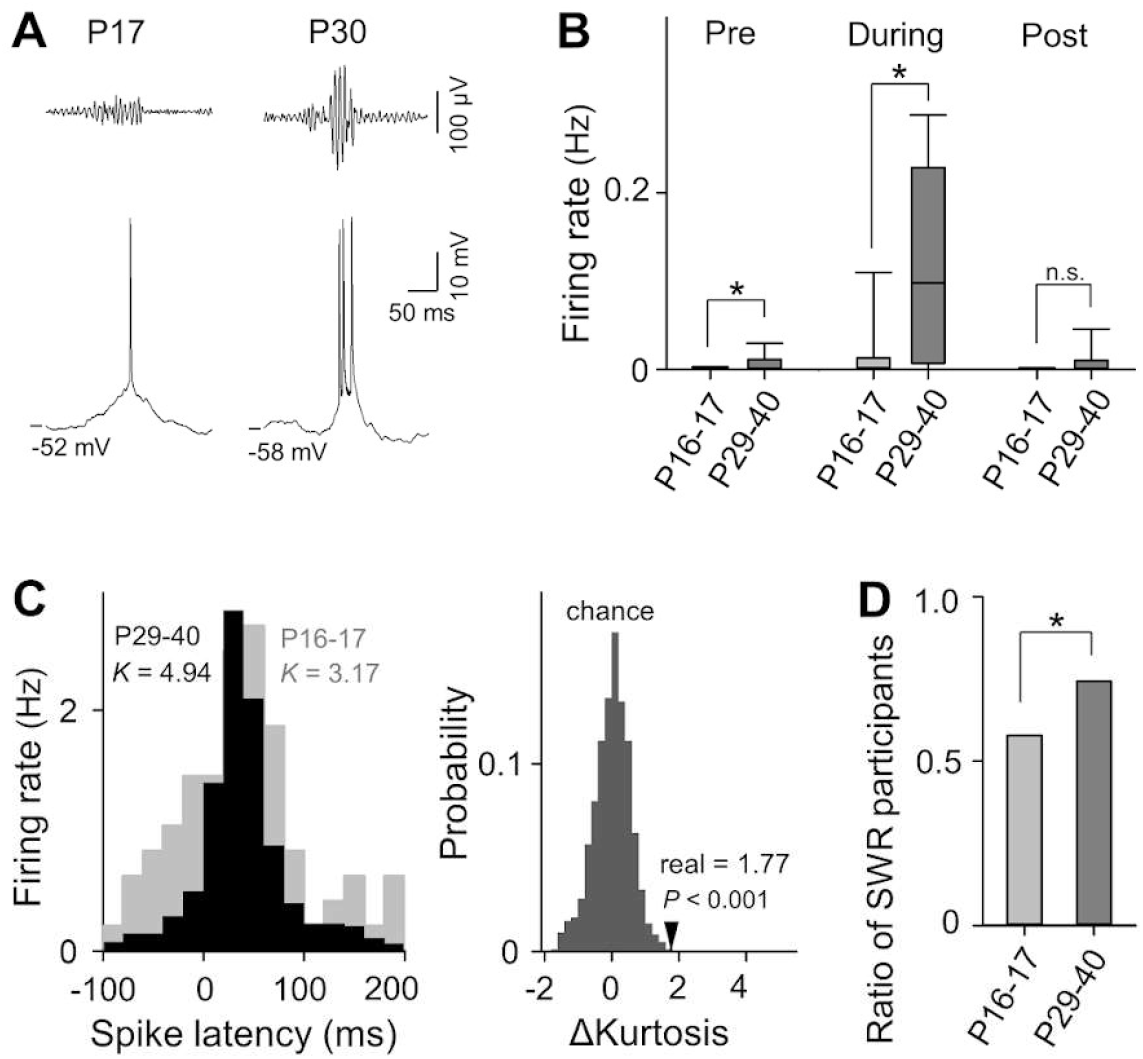
Comparison of firing activity between P16–17 and P29–40 mice. (**A**) Representative traces of bandpass-filtered LFPs (100-250 Hz) (top) and *V*m traces (bottom) during SWRs recorded from P17 and P30 mice. (**B**) Pre-, during- and post-SWR firing rates were all higher in P16–17 mice than in P29–40 mice. *P* = 0.033, 0.0053 and 0.16, *t*_63_ = -2.2, -2.9 and -1.4, Student’s *t*-test, for pre-, during- and post-SWR periods, respectively. *n* = 21 and 44 cells for P16–17 and P29–40 mice, respectively. (**C**) Left: The distributions of spike latencies relative to the SWR onsets in P16–17 mice (gray) showed higher kurtosis (*K* = 3.17) than that in P29–40 mice (black, *K* = 4.94). Right: The difference in the kurtosis (ΔKurtosis = 1.77) of the distributions of spike latencies between P16–17 and P29–40 mice (real), calculated from the left panel, was significantly larger than the chance level that was obtained from 1000 surrogates produced by randomly dividing the pooled spike latencies into two groups with the same numbers as the input data. (**D**) The proportion of cells that participated in SWRs to the total recorded cells was significantly higher in P29–40 mice than in P16–17 mice. *P* = 0.010, χ^2^ = 6.63, Chi-squared test.

We next quantified pyramidal cells that significantly increased their firing rates during SWRs. In each cell, the number of during-SWR spikes was compared to its null distribution, which was calculated from 1000 randomized surrogates in which spike times were randomly shifted within the recording period. Cells that had larger numbers of during-SWR spikes than the 95% confidence intervals of the null distributions were regarded as SWR participants. Of all recorded cells in P16–17 and P29–40 mice, 26/35 cells and 11/19 cells were SWR participants, the proportion of which was significantly higher in P29–40 mice (Figure 3D, *P* = 0.010, χ^2^ = 6.63, Chi-squared test). Therefore, premature CA1 pyramidal cells are more spontaneously active, but their spikes have less temporal selectivity to SWRs.

### Developmental increases in SWR-associated hyperpolarization

To explore mechanisms for the lack of temporal specificity of SWR-associated spikes in premature neurons, we focused on subthreshold *V*m dynamics (Valero et al., 2015; Hulse et al., 2016; Noguchi et al., 2022). First, all *V*m traces around SWR events were aligned to SWR onsets and averaged for cells during four developmental periods, *i*.*e*., P16–17, P21–22, P27–28 and P29–40 (Figure 4A). Consistent with a previous report (Hulse et al., 2016; Noguchi et al., 2022), the averaged trace in adult mice consisted of four components: (i) slow depolarization starting ∼1 s before SWRs, (ii) transient hyperpolarization ∼50 ms before SWR onset, (iii) large depolarization during SWRs, and (iv) hyperpolarization immediately after SWRs (Figure 4A bottom). Notably, the averaged *V*m trace of P16–17 mice lacked prominent hyperpolarizations both before and after SWRs, and the SWR-relevant depolarization had a longer duration than that of adult mice (Figure 4A top). The full widths at half maximum (FWHM) of depolarizations were smaller in older mice (Figure 4B, \**P* < 0.05, one-way analysis of variance (ANOVA) followed by Tukey‒Kramer *post hoc* test, *n* = 205, 516, 778 and 3782 SWRs for P16–17, P21–22, P27–28 and P29–40, respectively). The longer depolarizations in premature neurons may contribute to the lower selectivity of SWR-related spikes.

**Figure 4.**
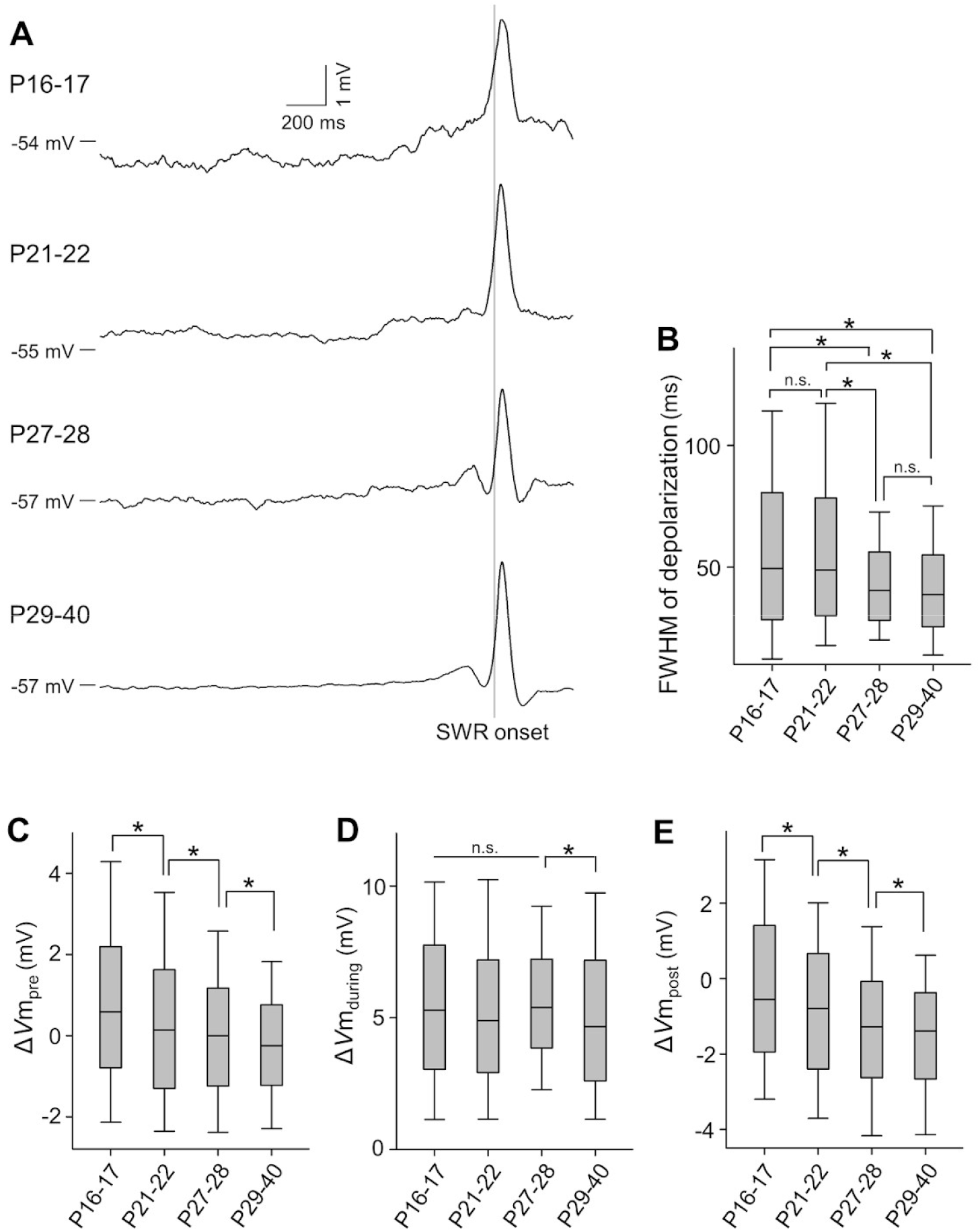
Developmental changes in peri-SWR *V*m dynamics. (**A**) For P16–17, P21– 22, P27–28 and P29–40, a total of 234, 593, 801 and 4138 *V*m traces in 21, 14, 9 and 44 cells from 15, 7, 5 and 20 mice were averaged relative to the SWR onsets (gray vertical lines), respectively. (**B**) Full widths at half maximum (FWHM) of depolarizations decreased during the fourth postnatal week. \**P* < 0.05, Tukey‒Kramer *post hoc* test after one-way ANOVA, *n* = 205, 516, 778 and 3782 SWRs for P16–17, P21–22, P27–28 and P29–40, respectively. (**C**) Δ*V*m_pre_ decreased with age throughout the developmental period. (**D**) Δ*V*m_during_ decreased after the fourth postnatal week. (**E**) Δ*V*m_post_ decreased with age throughout the developmental period. \**P* < 0.05, Tukey‒Kramer *post hoc* test after one-way ANOVA, *n* = 240, 595, 836 and 4159 SWRs in 21, 14, 9 and 44 cells from 15, 7, 5 and 20 mice for P16–17, P21–22, P27–28 and P29–40, respectively.

We quantified the *V*m changes (Δ*V*m) for the pre-, during- and post-SWR periods (Δ*V*m_pre_, Δ*V*m_during_, and Δ*V*m_post_, respectively). In the three parameters, Δ*V*m_pre_ and Δ*V*m_post_ gradually decreased with age, which indicates that more mature CA1 pyramidal cells received larger inhibitory inputs during the pre- and post-SWR periods (Figure 4C-E, \**P* < 0.05, one-way analysis of variance (ANOVA) followed by Tukey‒Kramer *post hoc* test, *n* = 240, 595, 836 and 4159 SWRs for P16–17, P21–22, P27–28 and P29–40, respectively). These results suggest that the maturation of peri-SWR inhibition restricts the temporal window for pyramidal cells to emit spikes during SWRs, which eventually results in temporally controlled activity in mature mice.

### Weak inhibitory and strong excitatory inputs in premature neurons

To directly examine the developmental changes in excitatory and inhibitory inputs, we voltage-clamped pyramidal cells at *V*ms of -70 and 10 mV and recorded excitatory and inhibitory postsynaptic conductances (EPSGs and IPSGs), respectively. The traces of EPSGs and IPSGs were averaged from 57 and 36 SWRs in 10 cells in P16–17 mice (Figure 5A). We plotted the time evolution of the mean values of peri-SWR conductances in the EPSG-versus-IPSG space (Figure 5A right) and found that EPSGs and IPSGs were linearly balanced, with IPSGs drifting slightly above EPSGs. In contrast, the mean traces of EPSGs and IPSGs for 54 and 62 SWRs in 7 cells from P29–32 mice demonstrate that IPSGs were dominant before SWR onset and remained greater than EPSGs throughout SWR events (Figure 5B), which is consistent with previous reports (Gan et al., 2017; Noguchi et al., 2022). The mean values of IPSGs during the pre-, during-, or post-SWR periods were calculated for the P16–17 and P29–32 datasets (Figure 5C). For all three periods, IPSGs were significantly smaller in P16–17 mice than in P29–32 mice (Figure 5C, *P* = 0.040, 2.4×10^−4^, 0.013, *D* = 0.28, 0.43 and 0.32 for the pre-, during- and post-SWR period, respectively, two-sample Kolmogorov‒Smirnov test, *n* = 36 and 62 SWR events for P16–17 and P29–40, respectively). In contrast, EPSGs were significantly larger in P16–17 mice than in P29–32 mice (Figure 5D, *P* = 2.7×10^−6^, 0.018, 0.0049, *D* = 0.48, 0.28 and 0.32 for the pre-, during- and post-SWR periods, respectively, two-sample Kolmogorov‒Smirnov test, *n* = 57 and 54 SWR events for P16–17 and P29–40, respectively). These results suggest that increasing inhibitory inputs and decreasing excitatory inputs shape the mature SWR-associated *V*m dynamics.

**Figure 5.**
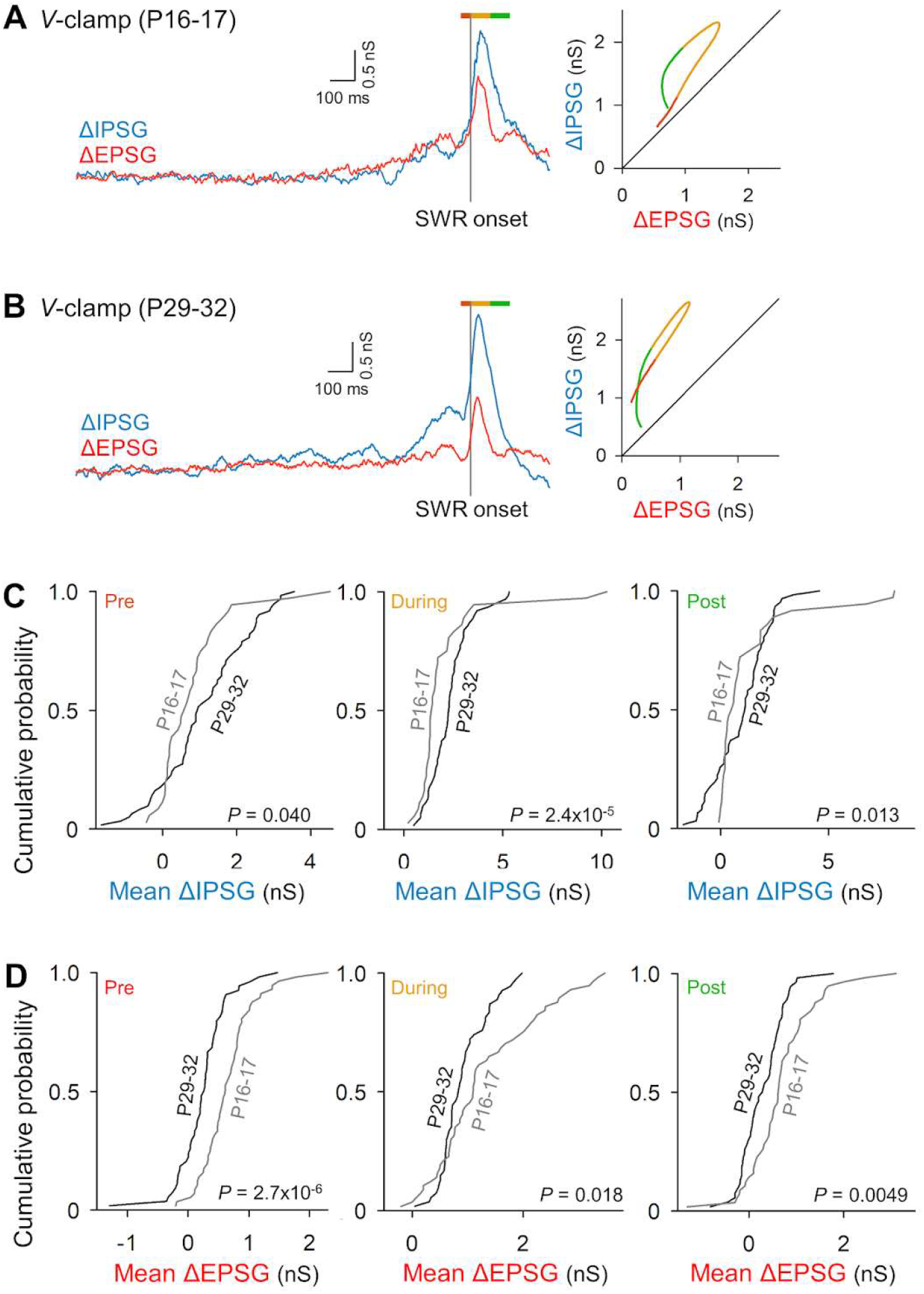
Developmental shifts to inhibitory dominance of peri-SWR synaptic inputs. Left: SWR onset-triggered average of the time changes in 57 EPSGs (red) and 36 IPSGs (blue) recorded from 10 cells in 7 P16–17 mice. The orange, yellow and green bars above the traces indicate pre-, during- and post-SWR periods, respectively. Right: The temporal coevolution of EPSGs and IPSGs in the left traces during the period shown by the colored bars in the EPSG-*vs*.-IPSG space. Pre-SWR EPSGs and IPSGs were balanced. The same as A, but for 54 EPSGs and 62 IPSGs recorded from 7 cells in 7 P29–40 mice. IPSGs increased earlier than EPSGs. (**C**) Cumulative probability distributions of mean values of pre- (left), during- (middle) and post- (right) SWR IPSGs in P16–17 mice (gray) and P29–40 mice (black). Mean IPSGs in P16–17 mice were smaller than those in P29–40 mice throughout the peri-SWR period. *D* = 0.28, 0.43 and 0.32, *P* = 0.040, 2.4×10^−4^, and 0.013 for the pre-, during- and post-SWR periods, respectively, two-sample Kolmogorov‒Smirnov test, *n* = 36 and 62 SWR events in 10 and 7 cells from 7 and 7 mice for P16–17 and P29–40, respectively. (**D**) The same as C, but for excitatory postsynaptic currents (EPSCs). Mean EPSGs in P16–17 mice were larger than those in P29–40 mice throughout the peri-SWR period. *P* = 2.7×10^−6^, 0.018 and 0.0049, *D* = 0.48, 0.28 and 0.32 for the pre-, during- and post-SWR periods, respectively, two-sample Kolmogorov‒Smirnov test, *n* = 57 and 54 SWR events in 10 and 7 cells from 7 and 7 mice for P16–17 and P29–40, respectively.

## Discussion

We discovered that hippocampal CA1 pyramidal cells at the beginning of SWR emergence exhibit high excitability with loose temporal specificity in both supra- and subthreshold activity. We also demonstrated gradual maturation of peri-SWR inhibition, which is thought to temporally control the spike times of pyramidal cells during SWRs in mature mice (English et al., 2014; Stark et al., 2015; Noguchi et al., 2022). The present study provides the first observation of *in vivo V*m dynamics of CA1 pyramidal neurons during postnatal development and proposes the neural mechanisms underlying less organized SWR-associated spike patterns in premature mice and their maturation through the third and fourth postnatal weeks (Farooq and Dragoi, 2019) at the single-cell *V*m level. Consistent with a previous report (Buhl and Buzsáki, 2005), we detected high-frequency oscillations at the end of the second postnatal week, but the frequencies in P14- 15 mice were low and cannot be considered mature SWRs. This observation may be attributable to the use of anesthesia in our experimental conditions (Ylinen et al., 1995). The increases in SWR event frequency and power are consistent with previous studies in freely moving rats (Buhl and Buzsáki, 2005; Farooq and Dragoi, 2019). In contrast to the previous study showing longer SWR events in older rats (Farooq and Dragoi, 2019), the duration of SWRs did not change, at least during the early developmental period we monitored. Considering that long-duration SWRs are linked to learning and memory (Fernández-Ruiz et al., 2019), we might have failed to capture the learning-dependent changes in SWR durations in anesthetized animals. In addition to the characteristics of individual SWRs, we reported an age-dependent increase in successive SWRs (Davidson et al., 2009; Yamamoto and Tonegawa, 2017), which are defined herein as more than two successive SWRs with interevent intervals of 250 ms. Successive SWRs are thought to arise from CA3 activity rather than from the entorhinal cortex, particularly under anesthesia (Yamamoto and Tonegawa, 2017). Successive SWRs have been linked to extended replays of long paths (Davidson et al., 2009). The developmental increase in successive SWRs may correspond to an increasing need for complex information processes, which are behaviorally accompanied by more active spatial explorations.

We described that CA1 pyramidal neurons in P16–17 mice had higher excitability (Chen et al., 2005), which was characterized by higher firing rates and more depolarized mean *V*m, than these neurons in adult mice. Previous *in vitro* studies have reported that the maturation of excitable membrane properties occurs during the first two postnatal weeks, and the resting *V*m of CA1 pyramidal cells becomes more hyperpolarized (Spigelman et al., 1992; Giglio and Storm, 2014), which seems incompatible with our data. The discrepancy may derive from the difference between *in vivo* and *in vitro* conditions; it should be noted that the anatomical structure is less preserved in slice preparations. We assume that the higher *V*m on P16–17 results from the immaturity of dendritic tonic inhibition (Cohen et al., 2000; Caraiscos et al., 2004; Ramos et al., 2004; Glykys and Mody, 2006; Groen et al., 2014) and that this effect is weakened in *in vitro* slice preparations because of the loss of synaptic connections from inhibitory interneurons to pyramidal cells. We also reported larger ISIs and fewer burst firings on P16–17 (Ranck, 1973; Harris et al., 2001). Spike bursts are usually initiated by synchronous excitatory inputs (Magee and Carruth, 1999). Indeed, more synchronous excitatory inputs from the CA3 subregion (Tyzio et al., 2003) and the entorhinal cortex (Donato et al., 2017) occurred in mature dendrites (Groen et al., 2014).

Frequent but temporally uncontrolled spikes cannot represent much information, which is the case in unorganized spike sequences in premature SWRs during postnatal development (Farooq and Dragoi, 2019). These loosely timed spikes may be due to broader depolarizations during SWRs in the premature hippocampus. In addition, we found that pre- and post-SWR inhibition, which emerges with age, narrowed the width of SWR-relevant depolarizations. This narrow depolarization may regulate the temporal window to allow pyramidal cells to fire spikes. Our voltage-clamp data demonstrated smaller inhibitory and larger excitatory inputs to pyramidal cells around SWRs in premature mice, which may underlie the age-dependent changes in peri-SWR *V*m dynamics. The smaller IPSGs are attributable to the immaturity of GABA_A_-receptor- mediated inhibition (Cohen et al., 2000; Groen et al., 2014) and presynaptic interneurons (Le Magueresse et al., 2011). We hypothesized that among the subtypes of CA1 interneurons, parvalbumin-positive basket cells mediated the growing inhibition based on their maturation in anatomical, intrinsic, and synaptic properties during the third and fourth postnatal weeks (Zhang et al., 1996; Doischer et al., 2008; Le Magueresse et al., 2011). This view is supported by the pivotal contribution of parvalbumin-positive basket cells to SWR-associated activity of pyramidal cells in adult mice (Lapray et al., 2012; Varga et al., 2014; Geiller et al., 2020). We also showed an age-dependent decrease in EPSGs, which appears inconsistent with the increasing excitatory synaptic transmission from CA3 to CA1 pyramidal cells throughout the developmental period (Hsia et al., 1998); however, because of the gradual increase in the inhibitory control of adult pyramidal cells in the CA1 and CA3 subregions (Romo-Parra et al., 2008; Sauer and Bartos, 2011; Groen et al., 2014) and the dominant inhibition during adult SWRs (Hájos et al., 2013; Hulse et al., 2016; Gan et al., 2017; Kajikawa et al., 2022), the increased inhibitory effect override the increased excitatory transmission.

It is notable that pre-SWR IPSGs were balanced with EPSGs, whereas post-SWR IPSGs already exceeded EPSGs on P16–17, characterizing the difference in synaptic activity between pre- and post-SWR periods at the beginning of the third postnatal week. CA1 pyramidal cells receive feedback inhibition during and after SWRs, particularly from parvalbumin-positive basket cells (Stark et al., 2014; Buzsáki, 2015), which shape during- and post-SWR hyperpolarization together with feedforward inhibition (Valero et al., 2015; Hulse et al., 2016). However, pre-SWR hyperpolarization reflect feedforward inhibition via CA1 interneurons that are driven by excitatory inputs from the CA2/3 subregions (Pouille et al., 2009; Bhatia et al., 2019; Noguchi et al., 2022). Thus, feedforward and feedback inhibition onto CA1 pyramidal cells could be different in developmental processes. Because the synaptic connectivity from CA3 to CA1 pyramidal cells continues to increase after the third postnatal week (Spigelman et al., 1992; Hsia et al., 1998), it is possible that the synaptic contacts from CA2/CA3 pyramidal cells to CA1 interneurons also develop during the third and fourth postnatal weeks.

In conclusion, our study presents SWR-associated *V*m dynamics *in vivo* and provides possible mechanisms underlying unorganized spike sequences of CA1 pyramidal cells in immature mice. The developmental time course of the maturation of *V*m dynamics, characterized by growing inhibition and its regulation of the time window for spikes, is compatible with the maturation of the sequential structure of replays (Farooq and Dragoi, 2019) and episodic memory (Travaglia et al., 2016). Several studies have suggested that instructive signals from the upstream network of the entorhinal cortex contribute to hippocampal replays (Schlesiger et al., 2015; Donato et al., 2017). In addition to this extrahippocampal mechanism, our results propose the concurrently maturing intrinsic and local-circuit properties in the hippocampus as factors for the developmental emergence of temporally organized activity during SWRs (Farooq and Dragoi, 2019).

## Acknowledgments

This work was supported by JST ERATO (JPMJER1801), the Institute for AI and Beyond of the University of Tokyo, and JSPS Grants-in-Aid for Scientific Research (18H05525, 20K15926).

## Data availability

The data that support the findings of this study are available from the corresponding author upon reasonable request.

## Code availability

The codes that support the findings of this study are available from the corresponding author upon reasonable request.

## Notes

**Conflicts of interest:** The authors declare no competing interests.

### Competing Interest Statement

The authors have declared no competing interest.

